# Road traffic delays and musculoskeletal health complaints among full-time bank employees: A cross-sectional study in Dhaka city

**DOI:** 10.1101/763052

**Authors:** Mohammad Ali, Gias U. Ahsan, Zakir Uddin, Ahmed Hossain

## Abstract

**Background:** The factors of road traffic delays (RTDs) have significant consequences for both commuters’ health and the country’s economy as a whole. Addressing the musculoskeletal health complaints (MHCs) among full-time employees has not been fully explored. The current study investigates the association between RTDs-related factors and MHCs among bank employees.

**Methods:** We conducted a cross-sectional study among full-time employees from 32 banks in Dhaka city. Descriptive statistics summarized the gaps in the socio-demographic and RTDs-related factors on the one-month prevalence of MHCs. Random intercept logistic regression models were used to identify the associate factors of the MHCs.

**Results:** Out of 628 full-time bank employees, the one-month prevalence of MHCs was 57.7%. The MHCs are more prevalent among adults of age group 40-60 years (68%) than the age group 20-40 years (54%). The one-month prevalence of lower back pain (LBP) was highest (36.6%), followed by neck pain (22.9%) and upper back pain (21.2%). Multilevel logistic regression analysis of employees showed that the odds of MHCs were lower among male employees (AOR=0.42, 95% CI= 0.27, 0.64), car commuters (AOR = 0.38, 95% CI=0.19-0.76, reference: bus commuters) and rickshaw commuters (AOR=. = 0.39, 95% CI=0.22-0.69, reference: bus commuters). The MHCs were significantly higher among employees with following factors: obesity (AOR= 1.50, 95% CI= 1.02-2.21), prolonged commute time to the office (AOR = 7.48, 95% CI =3.64-15.38) and working extended-time in a day (AOR= 1.50, 95% CI= 1.02-2.21).

**Conclusions:** The study indicates a high burden of musculoskeletal health complaints among the employees in Dhaka city, and the most prevalent complaint was low back pain. Our study suggests that factors related to road traffic delays might act synergistically on developing musculoskeletal problems in full-time employees.

## Background

Rising traffic congestion is an inescapable condition in large and growing cities across the world. Office commuters suffer the most of long delays due to peak-hour traffic congestion. Due to road traffic delays (RTDs), a substantial portion of working hours has to be left on the streets, which indirectly has an adverse impact on the economy. The delays force office commuters to adopt a long duration and the same posture in the vehicle for an extended period, which may induce musculoskeletal problems [1–3].

Most commuters globally suffer health problems due to RTDs. A longitudinal study in the United Kingdom demonstrated that longer commuting time is related to lower subjective health condition, lower health satisfaction, lower health status, poor sleep quality and higher body mass index [4]. Another study found longer commuting time related to worse mental health [5]. A study conducted in Norway found a high prevalence of subjective health complaints among long commuting railway workers [6]. Moreover, commuting long distance hurts physical activity, cardiorespiratory fitness, adiposity, and indicators of metabolic risk [7].

Commuters are also exposed to high concentrations of air pollutants and loud noises, and commuters in congested traffic suffer most of them. Prolonged and regular exposure to noise and air pollution can produce anxiety, depression, sleep disturbance, anger, and displeasure [8–10]. Commuters feel stressed and become anxious during the long commuting time [11–12]. This type of anxiety and depression due to longer commuting time could exacerbate musculoskeletal problems [13–14]. Despite a small increases in traffic volume, health risk is very high due to RTDs exposures, and it has a significant impact on public health [15–17].

Dhaka, the capital city of Bangladesh, is one of the most traffic-congested cities in the world. Office commuters in Dhaka city experience an inevitable overload on existing roads and transit systems every day. The first thing the commuters think of when it comes to congested roadways is the delay. The current road system cannot handle peak-hour loads without forcing many people to wait in line for limited road space, and it causes road traffic delays.

However, there is a research gap in estimating the effect of commuting time and musculoskeletal problems among full-time office commuters. Thus, the study has two objectives: (1) to estimate the prevalence of MHCs among bank employees of Dhaka city and (2) to find the association between factors related to RTDs and MHCs.

## Methods

### Study design and settings

This study was carried out from December 2018 to May 2019 among Bank employees in Dhaka city of Bangladesh. Dhaka is a large city where there are lots of commuters on the road, and most of the commuters experience heavy traffic congestion every day. According to UN World Urbanization prospects, more than twenty million people live in the Greater Dhaka area as of 2019 in approximately 300 square kilometers. There are more than 500,000 rickshaws, along with approximately 1.1 million registered motor vehicles operating every day in the city [18]. There are currently several hundreds of bank-branches available under 41 private and 9 public banks in Dhaka city. We followed the following eligibility criteria to include a participant in the study:

1. Full-time employees who worked in a bank for at least 1 year of Dhaka city corporation area;
2. We excluded nursing mothers and pregnant women, participants with chronic inflammatory arthritis (e.g., rheumatoid arthritis, gout, and ankylosing spondylitis);
3. Employees with disability were excluded;
4. Employees absent during the study period were excluded;
5. Older adults (60 and above) were excluded.

The eligible bank employees, who agreed to participate, received a questionnaire to complete either at the bank or home (collected it within 7 days from the bank). The completed questionnaires were returned, including the signed informed consent.

### Ethical consideration

The ethical committee of the Bangladesh University of Professionals (2019/273) and IRB of North South University (NSU-IRB-2019/54) approved the study. The objectives of the study, along with its procedure, was explained to the respondent, and written informed consent was taken from each respondent.

### Participants for analysis

We collected data conveniently from 83 bank-branches of 32 banks of Dhaka city. We provided a paper-based questionnaire to the 923 bank employees who met the eligibility criteria, and 652 of them returned the questionnaire. However, 628 participants completed the questionnaires, and we entered the data with an unknown id number for each participant in a personal computer for analysis.

## Study variables

### Measurement of musculoskeletal health complaints

The questions on musculoskeletal symptoms were based on the subscale of subjective health complaints by Eriksen et al. that measure health complaints experienced during last 30 days [19–20]. In this subscale of 8 items (shoulder pain, neck pain, lower back pain, upper back pain, arm pain, headache, leg pain during physical activity and Migraine) are included for which the severity of each complaint is scored on a four-point scale ranging from 0 (no complaint) to 3 (severe complaints). Thus, employees were asked to rate the occurrence of pain or discomfort using the 8 items with four answering categories (“no complaint”, “only once/a little”, “of short duration/ some”, “frequently/ serious”). Employees who answered, “no complaint”, “only once/a little”, “of short duration/ some” on all questions were classified as having no musculoskeletal health complaints. Those who answered “frequently/ serious” for one or more locations were classified as having musculoskeletal health complaints overall.

### Independent variables

Data regarding socio-demographic characteristics like age, gender, marital status, and in-house crowding were collected from the bank employees using a semi-structured questionnaire. The in-house crowding was defined as the total number of household members divided by total number of bedrooms in the house. We categorized the in-housing crowding in three groups: <=1.5, 1.5-2.0, and > 2.0. We collected data on the sleeping arrangement (firm/foam bed), smoking habit, physical activity. The physical activity was calculated based on METS scale [21]. We took into account occupational factors like duration of job experience in years, the average duration of daily working hours. We also collected data on chronic illness (diabetes and hypertension) from the employees.

### Road traffic delays related factors

Road traffic delays (RTDs) at a work zone include delays caused by deceleration of vehicles while approaching the work zone, reduced vehicle speed through the work zone, the time needed for vehicles to resume freeway speed after exiting the work zone, and vehicle queues formed at the work zone. In another study, RTD is defined as the unwanted journey time which can be calculated from the difference between the actual times required to commute and the time corresponding to the average speed of traffic under congested condition [22]. Thus, there are many factors involved in the calculation of delay equations. We took into account four factors to measure the RTDs: (1) average commute time for coming to office (minutes), (2) commuting distance (km) to office, (3) commuting mood and (4) overall subjective experience of traffic congestion (yes/no).

### Statistical analysis

We analyzed the data using the software, R 3.6.0. Descriptive statistics were calculated for all of the categorical variables (presented as frequencies and percentages). We performed a correlation analysis among the continuous independent variables to understand the multicollinearity among the data. We fitted a multilevel logistic model with the presence of MHCs. To run the multilevel logistic model, we chose a random intercepts model, with fixed slopes using the “glmer()” function from the “lme4” package of R. Here we specified the intercept varies by banks. The results are reported by the adjusted odds ratios (ORs), and corresponding p-values are also presented in a table. p-values less than 0.05 were considered statistically significant.

## Results

### Prevalence of MHCs

We present the one-month prevalence of MHCs among the 628 bank employees in Figure 1. Overall, 359 (57.2%) of the employees reported at least one musculoskeletal pain during the last month. Among the eight MHCs, the prevalence of lower back pain (LBP) was highest (36.6%), followed by neck pain (22.9%) and upper back pain (21.2%). Moreover, approximately 20% of the employees experienced headache, and 8.4% of the bank employees experienced migraine during the last one month.

**Figure 1:**
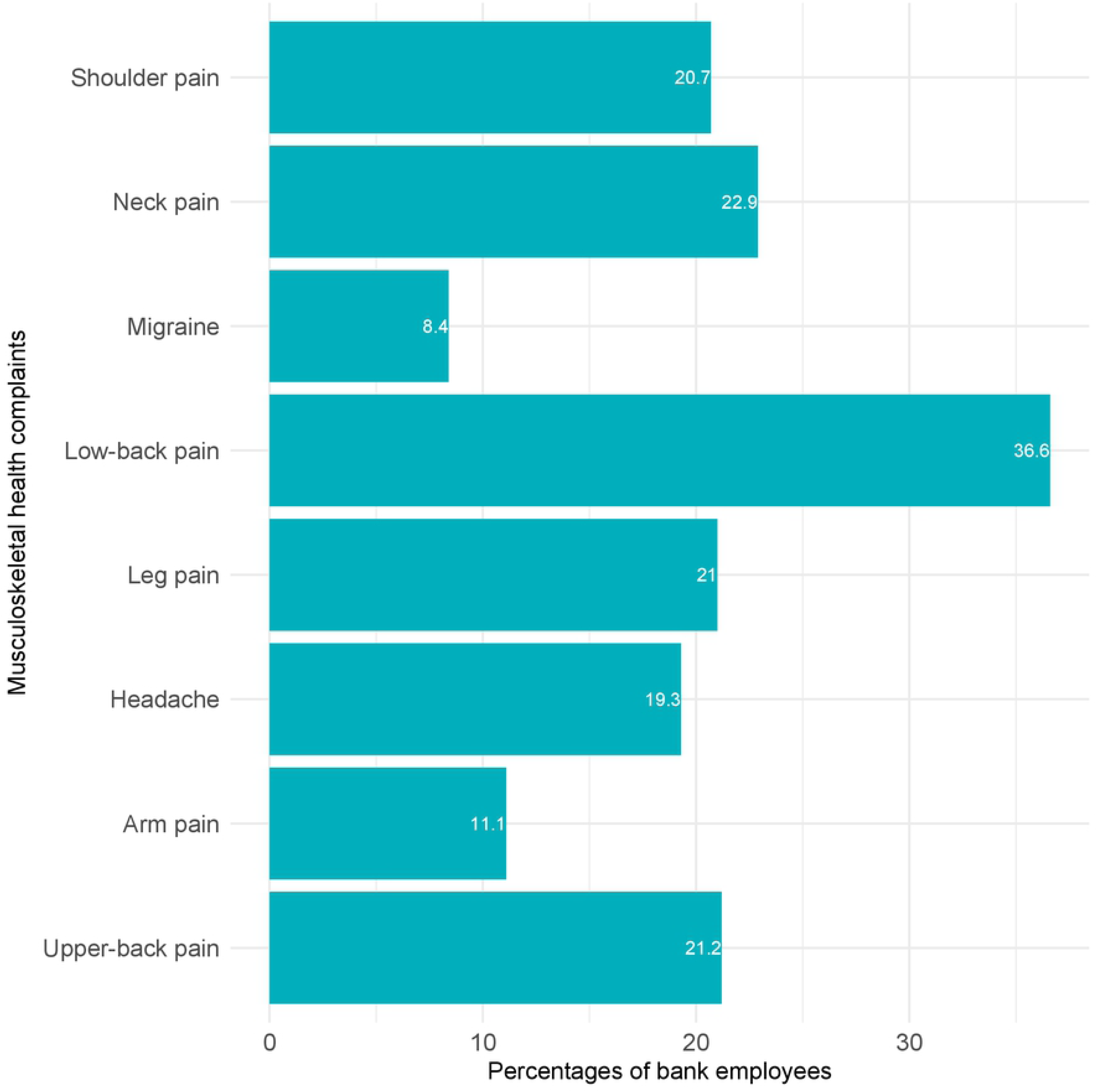
Percentage of bank employees corresponding to musculoskeletal health complaints.

### Multicollinearity analysis and Confounding bias

We discuss correlation analysis to quantify the association between two independent continuous variables to understand the multicollinearity of the data. Figure 2 shows the correlation plot among age, BMI, duration of the job (in a year), average commute time to the office (minutes), and commuting distance to the office (kilometers). The red dots present employees who usually experienced traffic congestion and blue dots present employees who did not experience traffic congestion. The figure shows the following four scenarios:

**Figure 2:**
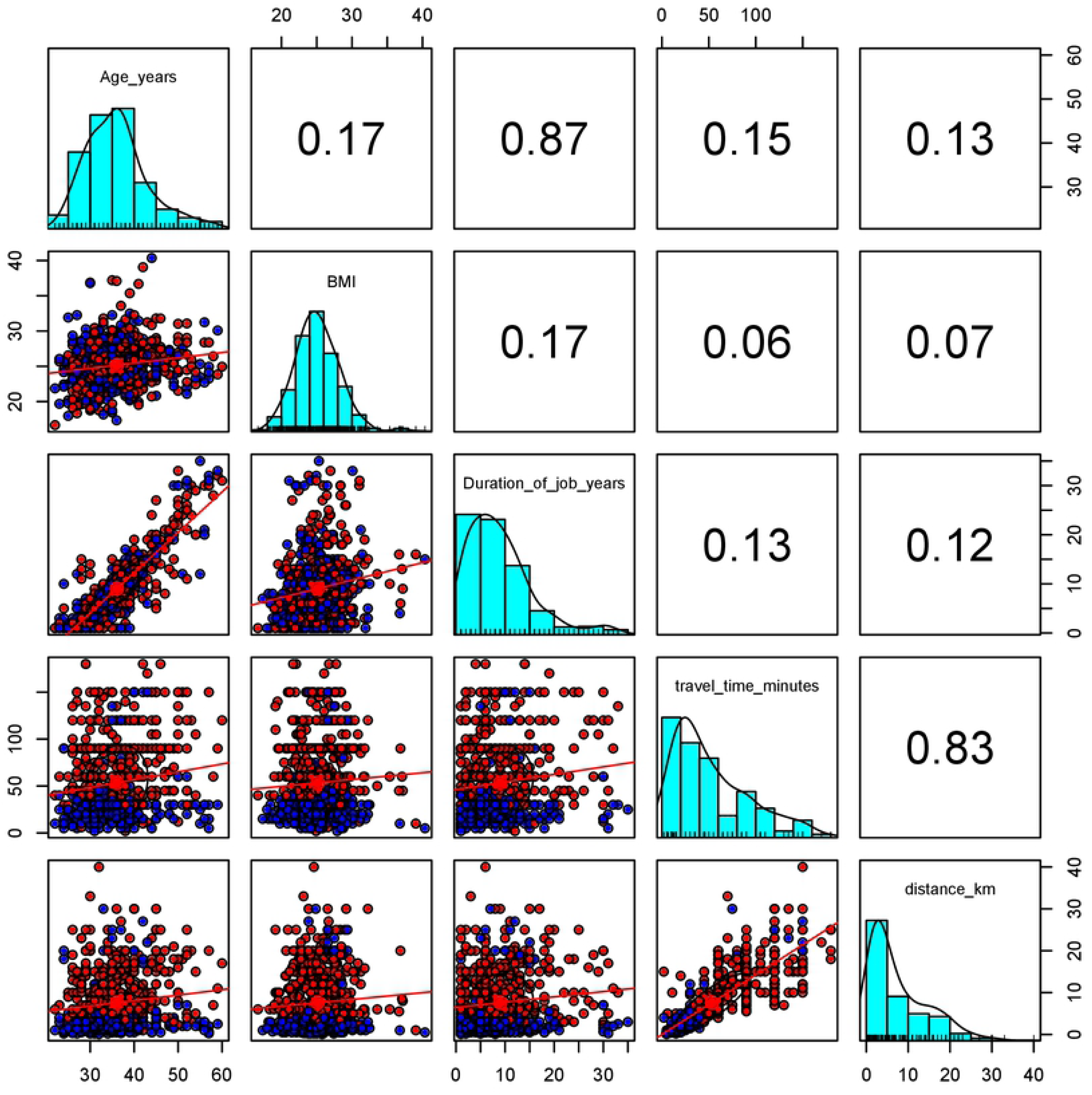
Correlation plot among age, BMI, duration of job (in year), commute time to office (minutes) and distance to office (in kilometer). The red dots present employees who usually experienced traffic congestion and blue dots present employees who do not experience traffic congestion.

1. It depicts a strong positive association (r=0.87) between employees age (years) and duration of the job (years).
2. It shows a weaker association between age and body mass index (r=0.17), and there is no association between BMI and average commute time (r approximately 0).
3. It depicts a strong positive association (r= 0.83) between average commute time and the commuting distance to the office.
4. It shows that blue dots are lying in the bottom of the correlation plot between average commute time and the commuting distance to the office. It indicates the employees with longer commute time, and long commuting distance to office reported traffic congestion.

The Venn-diagrams are presented in Figure 3 to investigate the relationship among age, BMI category, marital status, and musculoskeletal health complaints. The interior of the circle for the younger adults (age 20-40 years) in Figure 3 (left) represents the number of employees of the group, while the exterior represents some middle-aged adults (41-60 years) that are not members of the group. It appears from Figure 3 (left) that among the 392 married young adults, 56% of the participants reported MHCs. However, the Figure shows that 69% of the 124 married middle-aged adults reported MHCs. Therefore, the effect of age on the MHCs mixes with the effects of marital status for the MHCs that are present differentially by exposure status. Similar results are found with the factors of age and BMI category in Figure 3 (right). To avoid confounding bias, we kept only the BMI category in the final model.

**Figure 3:**
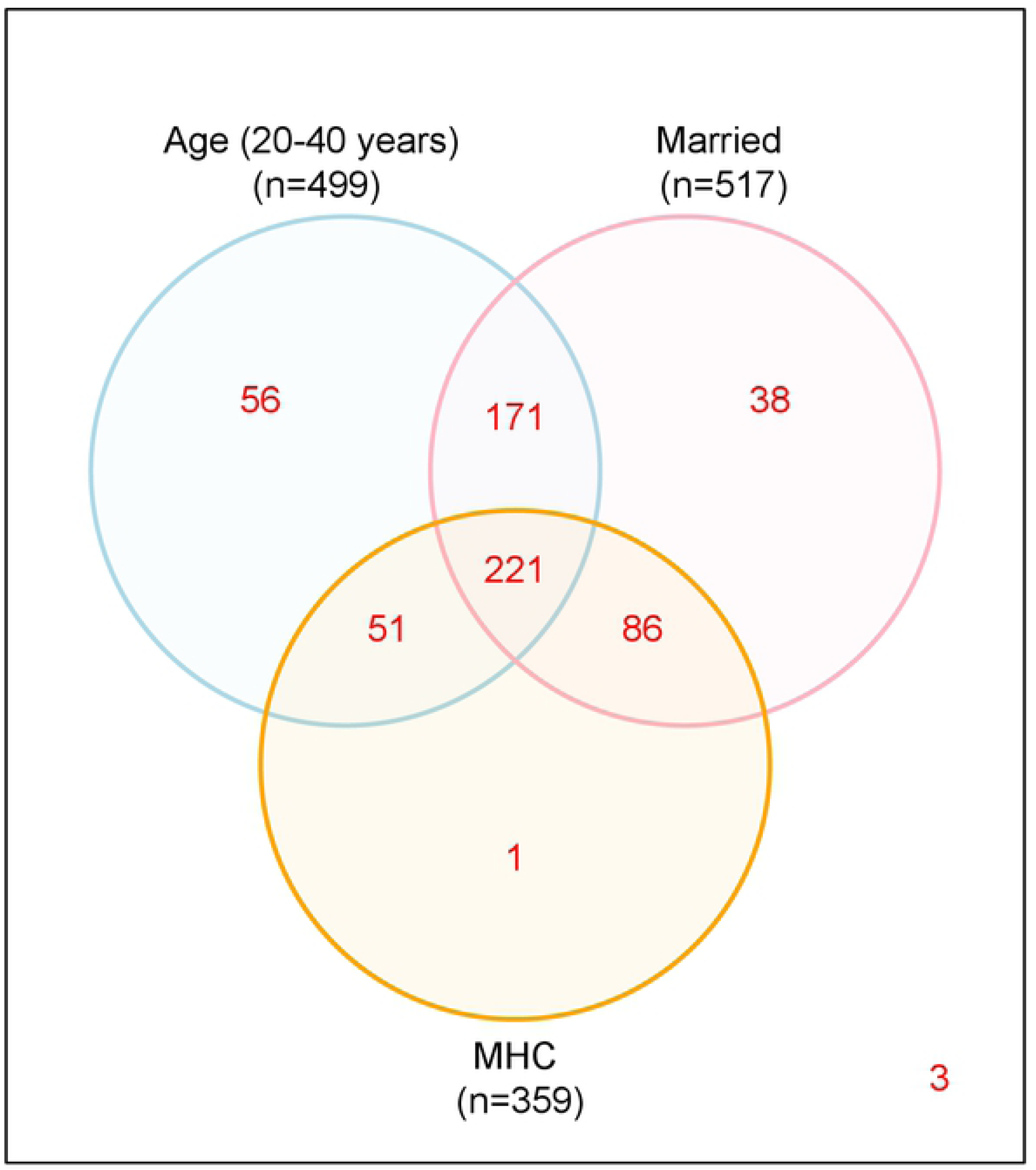
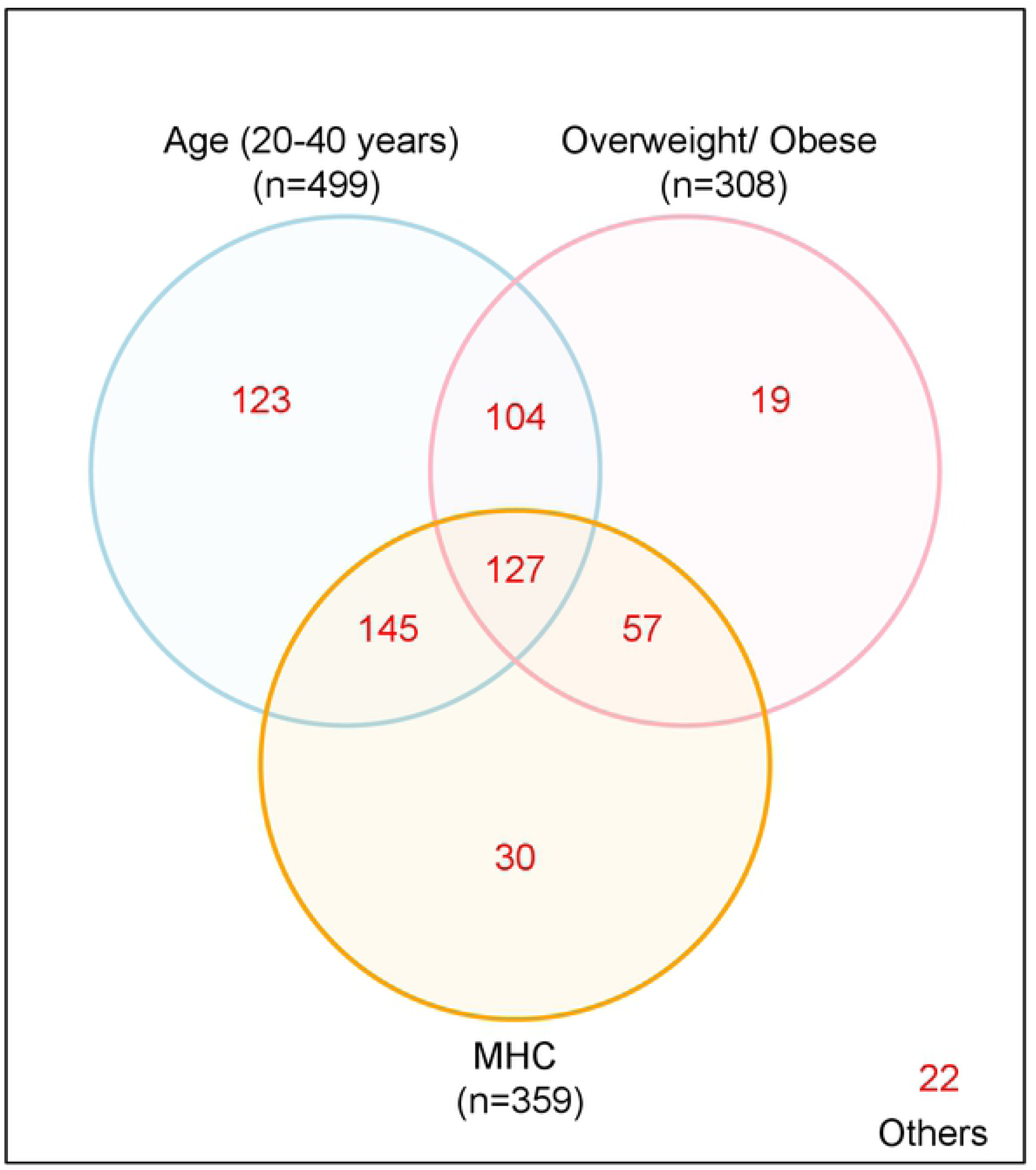
Venn-diagram of (left) Age, marital status and MHCs and (right) Age, Obesity and MHCs.

**Table 1:** Univariate Analysis: Socio-demographic and behavioral factors

| Factor | Categories | Musculoskeletal health complaints (MHCs) |  | Total (%) within categories | P-value* |
| --- | --- | --- | --- | --- | --- |
|  |  | Yes (Row %) | No |  |  |
| Age | 20-30 | 72 (51.8) | 67 | 139 (22.1%) | **0.044** |
|  | 31-40 | 200 (55.6) | 160 | 360 (57.2%) |  |
|  | 41-50 | 68 (68) | 32 | 100 (15.9%) |  |
|  | 50+ | 19 (67.9) | 9 | 28 (4.5%) |  |
| Gender | Male | 205 (54.9) | 168 | 373 (59.5%) | 0.185 |
|  | Female | 154 (60.6) | 100 | 254 (40.5%) |  |
| BMI | Normal | 175 (54.7) | 145 | 320 (51.0%) | 0.138 |
|  | Obese | 27 (71) | 11 | 38 (6.1%) |  |
|  | Overweight | 157 (58.4) | 112 | 269 (42.9%) |  |
| Marital Status | Married | 307 (59.5) | 209 | 516 (82.3%) | **0.019** |
|  | Unmarried | 52 (46.8) | 59 | 111 (17.7%) |  |
| Crowding | ≤ 1.5 | 185 (55.6) | 148 | 333 (53.1%) | 0.654 |
|  | 1.5-2 | 117 (59.4) | 80 | 197 (31.4%) |  |
|  | 2+ | 57 (58.8) | 40 | 97 (15.5%) |  |
| Sleeping mattresses | Firm bed | 302 (57.1) | 227 | 529 (84.4%) | 0.931 |
|  | Foam bed | 57 (58.2) | 41 | 98 (15.6%) |  |
| Smoking habit | No | 301 (58.3) | 215 | 516 (82.3%) | 0.285 |
|  | Yes | 58 (52.3) | 53 | 111 (17.7%) |  |
| Physical Activity | Sedentary | 115 (59.9) | 77 | 192 (30.6%) | 0.674 |
|  | Light | 213 (56.1) | 167 | 380 (60.6%) |  |
|  | Moderate to vigorous | 31 (56.4) | 24 | 55 (8.8%) |  |
| Diabetes/ Hypertension | No | 318 (56.9) | 241 | 559 (89.1%) | 0.685 |
|  | Yes | 41 (60.3) | 27 | 68 (10.9%) |  |

\* P-value is calculated from chi-square test.

### Univariate Analysis: Socio-demographic and behavioral factors

The mean ± SD age of the 628 participants was 36.1 ± 7 years. Most participants (80%) were young adults (20-40 years of age), and approximately 60% of the participants in the study were male. The young adults reported fewer MHCs (52%) compared to middle-aged adults (68%). The study observed a significant association between age groups and MHCs (p-value= 0.044). The majority of the participants 516 (82.3%) were married, and approximately 60% of the married employees reported about MHCs. The study illustrated a strong significant association between marital status and MHCs (p-value 0.019). In the study, female employees reported complaints more about musculoskeletal problems compared to male employees (61% versus 55%). Moreover, obese employees reported complaints (71%) more compared to the overweight (58.4%) or normal (54.7%) participants.

### Univariate Analysis: Job and road traffic delay related factors

The study finds all the road traffic delay related factors are univariately significant at a 5% significance level, and the results are given in Table 2. It appears that most of the employees used Bus (36%) to commute their office, and around 80% of them complain about musculoskeletal pains. Very few of the participants (13.4%) went to the office by walking or using a bicycle, and 33.3% of them had musculoskeletal pains. Table 2 also shows that as the duration of a job increases the MHCs increases. The employees who worked extended hours (>9 hours) regularly complained more about musculoskeletal pains. It appears that about 30% of employees reported commuting time for more than one hour, and 80% of them had musculoskeletal pains. Moreover, about half of the participants commute to offices by traveling more than 6 kilometers a day, and about 72% of them complained about musculoskeletal pains. Most of the participants (55%) experienced traffic congestion during the travel time to the office, and 82% of them complained about musculoskeletal pains.

**Table 2:**
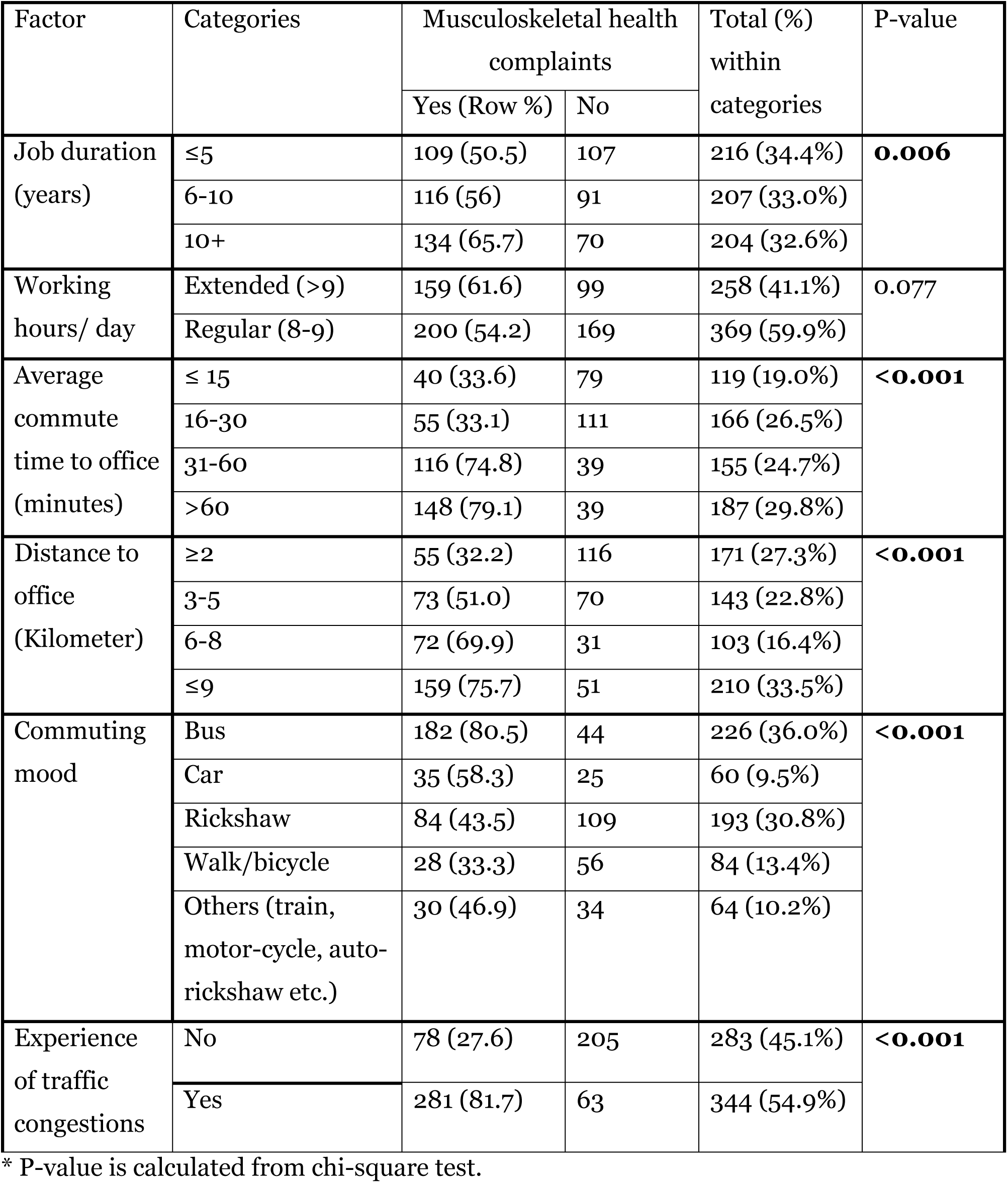
Univariate Analysis: Job and traffic congestion related factors

### Multilevel logistic regression Model

Table 3 shows the results of the adjusted odds ratio from two levels of logistic models with a random intercept for banks. We allow the intercept to vary randomly by each bank. The adjusted odds ratio (AOR) here is the conditional odds ratio for employees holding the factor constant as well as for employees with either the same bank or banks with identical random effects. We used two mixed logistic regression models to identify road traffic delay related factors that had significant associations with MHCs among young adults of age 20-40 years and all adults 20-60 years, respectively (Table 3). We consider the full model with the variables that were found significant with p-values <0.05 in the univariate analysis.

**Table 3:**
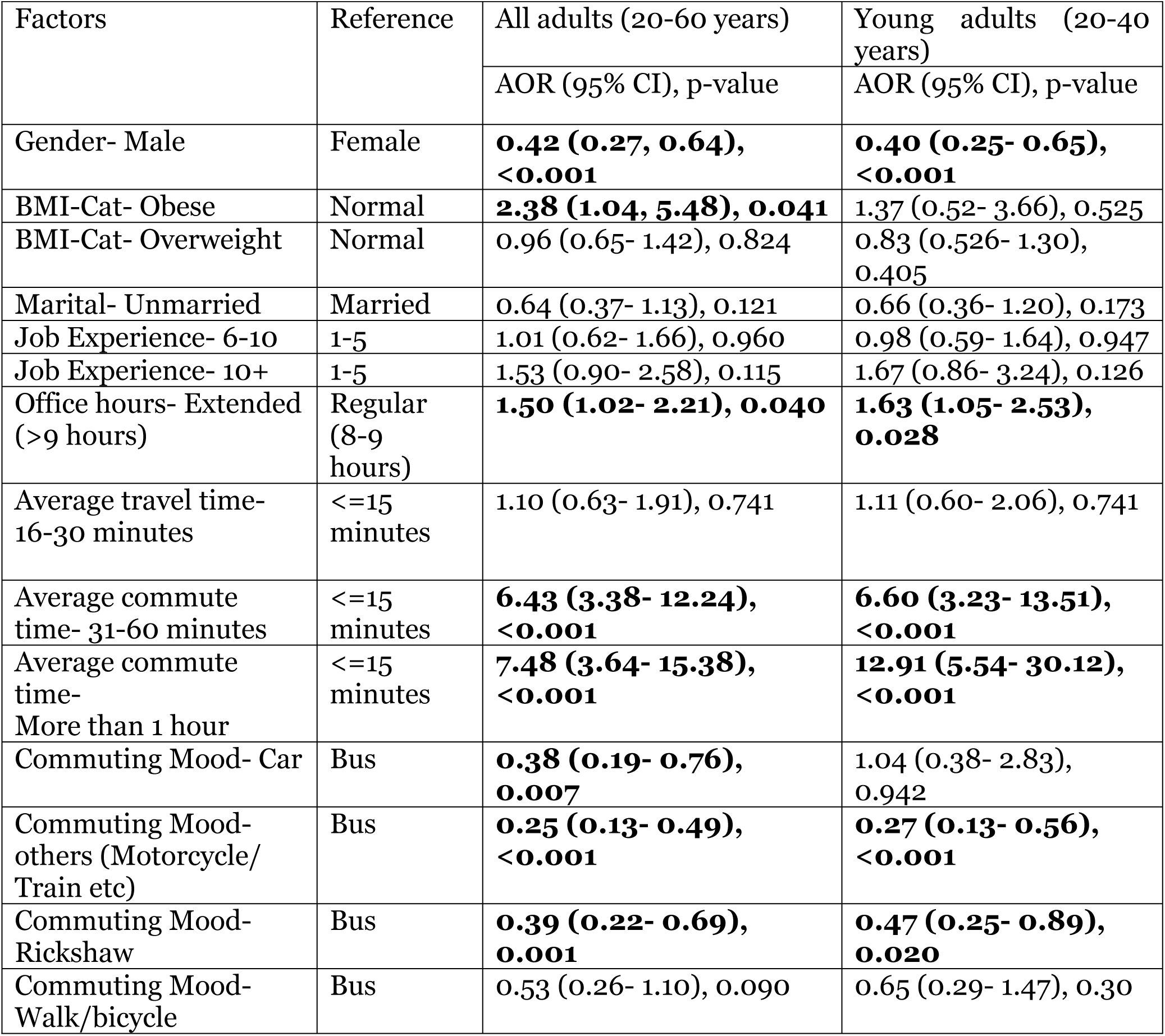
Result from multilevel Logistic model.

| Factors | Reference | All adults (20-60 years) | Young adults (20-40 years) |
| --- | --- | --- | --- |
|  |  | AOR (95% CI), p-value | AOR (95% CI), p-value |
| Gender- Male | Female | **0.42 (0.27, 0.64), <0.001** | **0.40 (0.25- 0.65), <0.001** |
| BMI-Cat- Obese | Normal | **2.38 (1.04, 5.48), 0.041** | 1.37 (0.52- 3.66), 0.525 |
| BMI-Cat- Overweight | Normal | 0.96 (0.65- 1.42), 0.824 | 0.83 (0.526- 1.30), 0.405 |
| Marital- Unmarried | Married | 0.64 (0.37- 1.13), 0.121 | 0.66 (0.36- 1.20), 0.173 |
| Job Experience- 6-10 | 1-5 | 1.01 (0.62- 1.66), 0.960 | 0.98 (0.59- 1.64), 0.947 |
| Job Experience- 10+ | 1-5 | 1.53 (0.90- 2.58), 0.115 | 1.67 (0.86- 3.24), 0.126 |
| Office hours- Extended (>9 hours) | Regular (8-9 hours) | **1.50 (1.02- 2.21), 0.040** | **1.63 (1.05- 2.53), 0.028** |
| Average travel time- 16-30 minutes | <=15 minutes | 1.10 (0.63- 1.91), 0.741 | 1.11 (0.60- 2.06), 0.741 |
| Average commute time- 31-60 minutes | <=15 minutes | **6.43 (3.38- 12.24), <0.001** | **6.60 (3.23- 13.51), <0.001** |
| Average commute time- More than 1 hour | <=15 minutes | **7.48 (3.64- 15.38), <0.001** | **12.91 (5.54- 30.12), <0.001** |
| Commuting Mood- Car | Bus | **0.38 (0.19- 0.76), 0.007** | 1.04 (0.38- 2.83), 0.942 |
| Commuting Mood- others (Motorcycle/ Train etc) | Bus | **0.25 (0.13- 0.49), <0.001** | **0.27 (0.13- 0.56), <0.001** |
| Commuting Mood- Rickshaw | Bus | **0.39 (0.22- 0.69), 0.001** | **0.47 (0.25- 0.89), 0.020** |
| Commuting Mood- Walk/bicycle | Bus | 0.53 (0.26- 1.10), 0.090 | 0.65 (0.29- 1.47), 0.30 |

### Employees aged 20-60 years

The results from mixed analysis indicate that male adults had lower odds of MHCs compared to the female adults (adjusted odds ratio, AOR= 0.40, 95% CI = 0.25-0.65). Overall, obesity was associated with an increased 1-month prevalence of MHCs. This association was significant for obesity (AOR= 2.38, 95% CI: 1.04, 5.48) regarding overall MHCs. It also appears from the analysis that the average commute time of more than 30 minutes to the office had a significant effect on the 1-month prevalence of MHCs (31-60 minutes AOR=6.43 and more than 1-hour AOR=7.48, respectively). In the analysis, employees using a car as the commuting mode can reduce 62% of the odds of 1-month prevalence of MHCs compared to the employees who used a bus (AOR=0.38, 95% CI=0.19-0.76).

### Employees aged 20-40 years

The following factors were significantly associated with an increased likelihood of musculoskeletal pains of the employees aged 20-40 years: female (AOR= 2.50, reference= male), extended office hours (AOR= 1.63, 95% CI= 1.05-2.53, reference: regular 8-9 office hours), average commute time to office-31-60 minutes (AOR=6.60, 95% CI=3.23-13.51, reference= <=15 minutes), average commute time to office-more than 1 hour (AOR=12.91, 95% CI=5.54-30.12, reference= <=15 minutes). Besides, employees who used rickshaw were 53% less likely to experience musculoskeletal pains than the employees who used a bus (AOR=0.47, 95% CI=0.25 – 0.89).

## Discussion

The study reveals that the one-month prevalence of MHCs among full-time bank employees is 57.2%. A study in China found that the three-months prevalence of musculoskeletal disorders (MSDs) among health care professionals was 68.3% [23]. Moreover, the one-year prevalence rate of MSDs among Iranian office workers was 52.8% [24] and among Thai office workers was 63% [25]. In Kuwait, 80% of bank employees experienced at least one episode of MSDs in the past year [26]. Another study in Saudi Arabia indicated that 56% of the adult population were in chronic musculoskeletal pain [27], which is similar to our findings.

In our study, the one-month prevalence of low back pain (LBP) was 36.6%, and neck pain was 22.9%. A study in Iran showed that one-year prevalence rate of LBP was 57.1% among office workers [28]. However, the prevalence of neck pain was leading MSD (60.2%) among Iranian office workers, which is much higher than our findings [28]. Moreover, research conducted among Malaysian commercial vehicle drivers suggested that among MSDs, LBP was the leading musculoskeletal problem with twelve-months prevalence of 60.4%, followed by neck pain (51.6%) and shoulder pain (35.4%) [29]. One-year prevalence of LBP among Israeli bus drivers was close to our findings (45.4%) [30].

However, two other studies conducted in Bangladesh found that the prevalence of LBP was 72.9% and 63% among nurses and garments workers, respectively [31–32].

Our study also investigated the associated factors for MHCs among full-time employees. The results of this study showed that age, gender, obesity, average duration of commuting time, distance to the office, and prolonged traffic congestion were significantly associated with MHCs among office commuters. We found the MHCs were more prevalent among female employees than male employees. A study conducted in Germany mentioned the point prevalence of back pain was 40% for women and 32% for men [33]. The existing literature is consistent as to whether obesity has an association with MHCs [34].

We found that traffic congestion and MHCs are positively associated. Similar findings were found previously in Norway, Malaysia, and UK [6, 30, 35]. There is also a strong positive association between prolonging commuting time to the office. A study described that long commuters encounter neck and lower back pain more frequently than short commuters [36]. Additionally, more recent research also found that the number and severity of MSDs were positively associated with long commuting time [6, 35]. Commuting long distance to the office is also positively associated with MHCs.

Furthermore, we did not find statistically significant associations between chronic diseases or physical activity and MHCs. However, chronic diseases in this study were assessed by asking employees about their current diabetes and hypertension condition, not by asking the duration of the diseases. A study showed that diabetes progression is positively associated with the presence of back pain [37]. However, we found a similar result of no association between physical activity and low back pain [38]. In our study, we did not ask about specific types of physical activity, but rather ascertained the number of hours of activity per week for all types of vigorous or moderate physical activity combined. More specifically, increasing amounts of exercise appear to have a variable effect on musculoskeletal health, but specific categories of exercise may affect musculoskeletal health negatively.

There are many strengths to this study. Only a few prior studies have described gaps in MHCs among full-time employees. Our study is the first to highlight the association between factors related to RTDs and MHCs. The study participants were bank employees who were homogenous because of their job nature, sitting arrangement, and the environment of the office was almost the same. Limitations include the self-reported data collected through study participant questionnaires. Owing to the cross-sectional study design, MHCs differences observed in this paper cannot be interpreted as causative. In a future goal, a multi-professional study may help determine the overall generalizability of our results to the full-time employee as a whole.

## Conclusions

The study finds a high burden of musculoskeletal problems among full-time bank employees in Dhaka city, and the most prevalent complaint was low back pain. Traffic congestion is inevitable in the city, and it has negative consequences on musculoskeletal problems. Long-distance commuters and prolonged commuters by bus are more likely to report musculoskeletal pains. Bus commuters can break up the continuous sitting and overcome the deleterious effects of road traffic delays. Whether full-time employees may be able to improve the musculoskeletal problems by reducing commute time and losing weight must be further investigated.

## Consent to Publish

Not applicable.

## Availability of data

Click here for the data file http://individual.utoronto.ca/ahmed_3/index_files/data/data.html.

### Competing interests

The authors declare that they have no competing interests.

## Funding

A small seed fund was available from Bangladesh University of Professionals to conduct the study.

## Author’s contributions

MA and AH participated in study conception, design and coordination of the manuscript. GUA, and ZU reviewed the manuscript and helped to draft the manuscript. AH also performed statistical analysis and helped to draft the manuscript. All authors approved the final manuscript.

## Acknowledgements

All the authors acknowledge the participants for providing us the information to conduct the study. We also thank Professor Norman K. Swazo for assistance with English language usage, grammar, and spelling that greatly improved the manuscript.

